# Dual role of Vascular Endothelial Growth Factor-C (VEGF-C) in post-stroke recovery

**DOI:** 10.1101/2023.08.30.555144

**Authors:** Yun Hwa Choi, Martin Hsu, Collin Laaker, Melinda Herbath, Heeyoon Yang, Peter Cismaru, Alexis M. Johnson, Bailey Spellman, Kelsey Wigand, Matyas Sandor, Zsuzsanna Fabry

**Affiliations:** Department of Medicine, University of Wisconsin-Madison, Madison, WI 53705, USA; Waisman Center, University of Wisconsin-Madison, Madison, WI 53705, USA; Department of Microbiology and Immunology, University of North Carolina, Chapel Hill, NC 27599, USA; Neuroscience Training Program, University of Wisconsin-Madison, Madison, WI 53705, USA; Department of Pathology and Laboratory Medicine, University of Wisconsin-Madison, Madison, WI 53705, USA; College of Agricultural and Life Sciences, University of Wisconsin-Madison, Madison, WI 53705, USA

## Abstract

Using a mouse model of ischemic stroke, this study characterizes stroke-induced lymphangiogenesis at the cribriform plate (CP). While blocking CP lymphangiogenesis with a VEGFR-3 inhibitor improves stroke outcome, administration of VEGF-C induced larger brain infarcts.

**Abstract:** Cerebrospinal fluid (CSF), antigens, and antigen-presenting cells drain from the central nervous system (CNS) into lymphatic vessels near the cribriform plate and dural meningeal lymphatics. However, the pathological roles of these lymphatic vessels surrounding the CNS during stroke are not well understood. Using a mouse model of ischemic stroke, transient middle cerebral artery occlusion (tMCAO), we show that stroke induces lymphangiogenesis near the cribriform plate. Interestingly, lymphangiogenesis is restricted to lymphatic vessels at the cribriform plate and downstream cervical lymph nodes, without affecting the conserved network of lymphatic vessels in the dura. Cribriform plate lymphangiogenesis peaks at day 7 and regresses by day 14 following tMCAO and is regulated by VEGF-C/VEGFR-3. These newly developed lymphangiogenic vessels transport CSF and immune cells to the cervical lymph nodes. Inhibition of VEGF-C/VEGFR-3 signaling using a blocker of VEGFR-3 prevented lymphangiogenesis and led to improved stroke outcomes at earlier time points but had no effects at later time points following stroke. Administration of VEGF-C after tMCAO did not further increase post-stroke lymphangiogenesis, but instead induced larger brain infarcts. The differential roles for VEGFR-3 inhibition and VEGF-C in regulating stroke pathology call into question recent suggestions to use VEGF-C therapeutically for stroke.

## Introduction

Stroke is the fifth leading cause of death and the leading cause of long-term disability according to the American Heart Association (Virani et al., 2020). Currently, there is only one FDA-approved drug available for ischemic stroke patients, tissue plasminogen activator (tPA) (Kim, 2019; Özlüer and Avcil, 2017; Knecht et al., 2018). However, tPA is only given to about 2-6% of ischemic stroke patients due to its narrow therapeutic window and its potential risk of adverse effects such as intracerebral hemorrhagic conversion (Kim, 2019; Knecht et al., 2018; Alberts, 2017; Gravanis and Tsirka, 2008; Knecht et al., 2017; Lin et al., 2018; Barber et al., 2001; Miller et al., 2011). For years, researchers have tried to develop better therapeutic approaches for ischemic stroke, but most clinical trials have fallen short (Xu and Pan, 2013; Chen and Wang, 2016; Cheng et al., 2004; Stroke Therapy Academic Industry Roundtable II (STAIR-II), 2001).

Stroke leads to acute brain edema that contributes to blood-brain barrier disruption, inflammation, and homeostatic disbalance (Rosenberg, 1999). Resolution of brain edema has been associated with functional lymphatic clearance of the brain (Si et al., 2006; Chen et al., 2019; Hu et al., 2020). Disparagingly, both inducing lymphatic vessel formation or inhibiting lymphangiogenesis have been proposed as potential therapies for the resolution of post-stroke brain edemas and accelerating post-stroke recovery (Esposito et al., 2019; Simoes Braga Boisserand et al., 2023; Kim et al., 2021). One of the difficulties of designing lymphatic targeting therapies in stroke lies in the uncertainty of the targeted pathways or exact mechanisms. Different lymphatic vessels around the brain could mediate lymphatic clearance, including the dural meningeal lymphatics (Da Mesquita et al., 2018; Wen et al., 2018), basal lymphatics (Ahn et al., 2019), and the cribriform plate lymphatics (Hsu et al., 2019). These lymphatic vessels have been shown to be involved in transporting fluid, waste, and immune cells in a variety of pathological states (Alitalo, 2011; Ma et al., 2017). Previously, our lab has shown that lymphatic vessels near the cribriform plate proliferate and expand in experimental autoimmune encephalomyelitis (EAE), a mouse model of autoimmune-mediated neuroinflammation. These newly formed lymphatic vessels have immunoregulatory functions that may manage neuroinflammatory diseases (Hsu et al., 2019, 2022). Intriguingly, multiple labs reported that stroke induces lymphangiogenesis at the dural meningeal lymphatics in mouse intracerebral hemorrhage (ICH) (Tsai et al., 2022) and photothrombosis mouse stroke models (Yanev et al., 2020), however, whether targeting these pathways would be beneficial for stroke recovery is still unknown.

Manipulating the VEGFR-3/VEGF-C pathway has been proposed as a potential new therapy for stroke treatment (Simoes Braga Boisserand et al., 2023; Tsai et al., 2022; Yanev et al., 2020; Esposito et al., 2019). VEGF-C is commonly secreted by immune cells during inflammatory events and binds to VEGFR-3 expressed on lymphatic vessels to stimulate lymphangiogenesis (Flister et al., 2010) . VEGF-C has been documented to increase in the CNS in days following stroke (Gu et al., 2001; Shin et al., 2008; Bain et al., 2013). However, the contribution of VEGFR-3/VEGF-C signaling to stroke pathology still remains controversial. Some reports indicate VEGF-C may improve post-stroke recovery by promoting the survival of neural stem cells (Matta et al., 2021) or promoting increases in CNS clearance through expansion of dural meningeal lymphatic vessels (Tsai et al., 2022). Conversely, others have suggested that early widespread blockade of VEGFR-3 and VEGF-C signaling can reduce ischemic infarction size through reductions in pro-inflammatory immune activation within the cervical lymph nodes (Esposito et al., 2019).

To clarify the role of lymphatics in post-stroke pathology, we investigated the impact of ischemic stroke in multiple CNS – associated lymphatic regions (cribriform plate, dura, and cervical lymph nodes) and test the effects of VEGFR-3/VEGF-C signaling on stroke outcome. Using a transient middle cerebral artery occlusion (tMCAO) mouse model, we found that after 3 days of tMCAO, lymphangiogenesis occurs near the cribriform plate which peaked at day 7 and decreased by day 14. Lymphatic expansion was associated with increased immune cell interaction, particularly with macrophages and dendritic cells. As expected, lymphangiogenesis near the cribriform plate appeared to occur via increased VEGFR-3/VEGF-C signaling. The presence of VEGF-C secreting CCR7^+^, CD11b^+^, CD11c^+^ migratory dendritic cells near the cribriform plate lymphatics indicate that these cells are recruited to this area and are the most likely sources of VEGF-C. Lymphangiogenic vessels near the cribriform plate transported fluids from the CNS after 7 days of tMCAO. In agreement with this, downstream CNS-draining cervical lymph nodes (but not other lymph nodes) also experienced post-stroke lymphangiogenesis. In contrast to our findings for lymphangiogenesis near the cribriform plate and cervical lymph nodes, we did not see evidence for this at the cranial dual lymphatic vasculature. Cranial dural meningeal lymphatics adapted to stroke-induced neuroinflammation by dilation 3 days following tMCAO. Confirming the therapeutic findings by others (Esposito et al., 2019), inhibition of VEGFR-3 signaling not only blocked lymphangiogenesis near the cribriform plate and dilation of cranial dural LVs, but also led to smaller brain infarcts and improved motor ability following tMCAO. With the goal of increasing stroke neuroinflammation-induced drainage of the brain, we administered VEGF-C156 through the left common carotid artery after tMCAO. However, post-stroke VEGF-C treatment induced larger brain infarcts without further inducing changes on lymphatics in the cranial dura and cribriform plate lymphatics. Our data support that VEGFR-3-dependent lymphangiogenesis near the cribriform plate is induced after ischemic stroke in mice but also show that manipulation of VEGF-C/VEGFR-3 interaction may not lead to better therapeutics for stroke.

## Results

### Lymphatic vessels near the cribriform plate, not at the dorsal cranial region, undergo lymphangiogenesis after tMCAO

In order to assess the role of lymphatics in the CNS after stroke in mice, we first looked to confirm if lymphangiogenesis occurs near the cribriform plate after tMCAO. Our lab previously showed that lymphangiogenesis occurs at the cribriform plate in the EAE model of autoimmune mediated neuroinflammation (Hsu et al., 2019). After inducing tMCAO for 60 minutes in wild-type male mice, stroke lesion size was measured using T2-weighted MRI and then mice were sacrificed on day 3, 7, or 14. Whole heads were fixed and decalcified for immunohistochemistry (IHC). Brain infarction was confirmed at all time points in sham and tMCAO groups. Infarction areas were calculated to be between 30-40% across day 3, 7, and 14 with slight lower percentages at day 14 after tMCAO (Fig. 1A). We then sectioned the cribriform plate-olfactory region, and stained it with Lyve-1-fluorescent antibody to identify lymphatic vessels at each time point. Average Lyve-1^+^ vessel volumes were significantly elevated at day 3, peaking at day 7, and then decreased at day 14 in the tMCAO group, while the sham group did not show changes across those time points (Fig. 1B). Additionally, we quantified the number of loops and sprouts to confirm lymphangiogenesis near the cribriform plate. Loops and sprouts morphology is a signal of increased complexity and can be used to estimate lymphatic proliferation (Kajiya et al., 2009). Number of loops was increased at the cribriform plate among tMCAO groups at both day 3 and 7, while the number of sprouts was increased at day 3, 7, and 14 (Sup. Fig. 1A). We also measured the length of lymphatic vessels near the cribriform plate starting from nasal mucosa to the tip of lymphatic vessels between the olfactory bulbs. The lengths were increased at day 3 and 7, and then decreased to similar length of sham groups at day 14 (Sup. Fig. 1B). These data indicate that lymphangiogenesis near the cribriform plate occurs and initiates around day 3 and regresses by day 14 after tMCAO.

**Figure 1.**
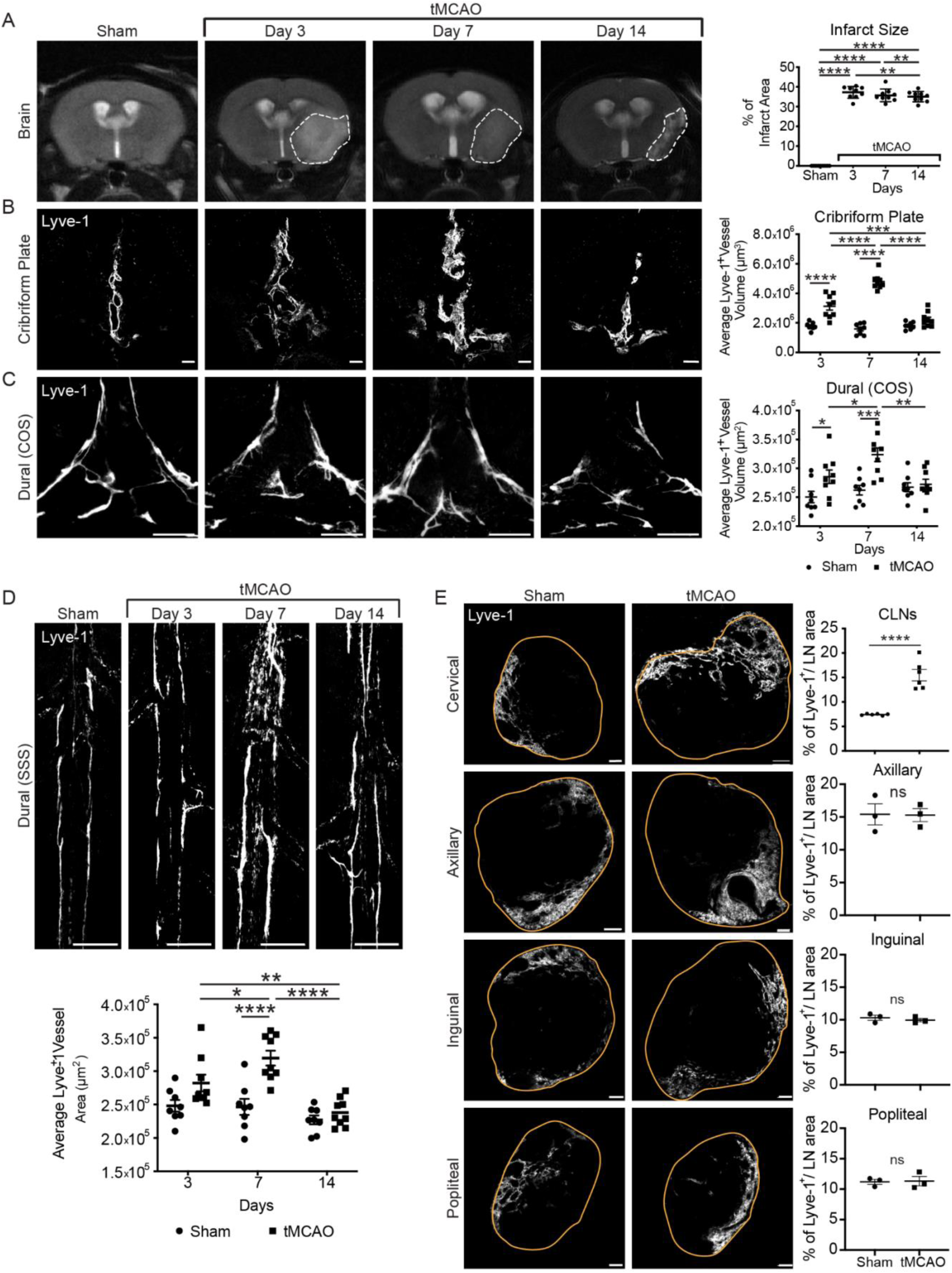
Cribriform plate lymphatic vessels undergo lymphangiogenesis following tMCAO. **(A):** T2-weighted MRI images of brain sections between sham and tMCAO at day 3, 7, and 14. Representative images of MRI brain scans after sham and tMCAO. White dashed lines mark brain infarcts which are quantified into a graph (n=8 mice for sham, n=9 mice for tMCAO; mean ± SEM, *p ≤ 0.05, ****p ≤ 0.0001, two-way ANOVA) **(B):** Coronal sections of cribriform plate areas were immunolabeled for Lyve-1 to visualize lymphatic vessels after 3, 7, and 14 days of sham or tMCAO. Representative confocal images of lymphatic vessels near the cribriform plate. Scale bars = 100 µm. Quantitation of average Lyve-1^+^ vessel volumes near the cribriform plate (n=8 mice for sham, n=9 mice for tMCAO; mean ± SEM, *p ≤ 0.05, **p ≤ 0.01, ***p ≤ 0.001, ****p ≤ 0.0001, two-way ANOVA). **(C):** Confluence of sinuses (COS) of dural lymphatic vessels were stained with Lyve-1 antibody and imaged using confocal microscopy after sham and tMCAO at day 3, 7, and 14. Representative confocal images of COS. Scale bars = 500 µm. Quantitation of average Lyve-1^+^ vessel areas in the COS (n=8 mice for sham, n=9 mice for tMCAO; mean ± SEM, *p ≤ 0.05, **p ≤ 0.01, ***p ≤ 0.001, ****p ≤ 0.0001, two-way ANOVA). **(D):** Superior sagittal sinus (SSS) of dural lymphatic vessels were immunolabeled with Lyve-1 fluorescent antibody and imaged after sham and tMCAO at day 3, 7, and 14. Representative confocal images of SSS. Scale bars = 500 µm. Quantitation of average Lyve-1^+^ vessel areas in the SSS (n=8 mice for sham, n=9 mice for tMCAO; mean ± SEM, *p ≤ 0.05, **p ≤ 0.01, ***p ≤ 0.001, ****p ≤ 0.0001, two-way ANOVA). **(E):** Sections of cervical lymph nodes and peripheral lymph nodes such as axillary, inguinal, and popliteal were immunolabeled with Lyve-1 fluorescent antibody after 7 days of sham and tMCAO. Representative confocal images of each lymph node with orange lines to define the areas of lymph nodes. Scale bars = 100 µm. Quantitation of percentage of Lyve-1^+^ vessel area over lymph node areas (n=6 mice per group for cervical lymph nodes, n=3 mice per group for axillary, inguinal, and popliteal lymph nodes; mean ± SEM, ****p ≤ 0.0001, unpaired Student’s t-test).

We tested whether the cranial dural meningeal lymphatics would undergo changes after tMCAO. Similar to lymphatics near the cribriform plate, average Lyve-1^+^ vessel areas of the confluence of sinuses (COS) of dural lymphatics were increased at day 3 and 7, and then decreased at day 14 after tMCAO compared to sham groups (Fig. 1C). When we looked at the superior sagittal sinus (SSS) of the dural lymphatics, there was an increase in Lyve-1^+^ vessel areas at day 7 which decreased by day 14 after tMCAO (Fig. 1D). We quantified the number of loops and sprouts of both COS and SSS in dural meningeal lymphatics, but there were no differences in both areas between sham and tMCAO groups. Quantitation of lymphatic diameter in the COS showed increases at day 3 and 7, and regressed back to a similar diameter of sham groups at day 14 (Sup. Fig. 2C). Diameter of vessels in the SSS showed increases at day 7 of tMCAO compared to sham (Sup. Fig. 2B). Together this data indicates that cranial dural meningeal lymphatics undergo lymphatic vessel dilation, but may not necessarily undergo lymphangiogenesis as seen in the cribriform plate.

We next tested whether lymphangiogenesis expanded to the peripheral lymph nodes following stroke. Here increased Lyve-1^+^ areas were observed in cervical lymph nodes (CLNs) of tMCAO 7 days following tMCAO, however, these changes were specific to the CLNs, since other peripheral lymph nodes such as axillary, inguinal, and popliteal had no changes in Lyve-1 expression profiles during the observation period (Fig. 1E). These data support the strong regional cooperation of adaptive lymphatic responses restricted to the cervical lymphatics and the CP lymphatic vasculature; further supporting the unique nature of cribriform plate lymphatic vessels and their similarities to peripheral lymphatic vessels.

### Lymphangiogenic vessels in the CP can retain immune cells and drain fluid after tMCAO

To test whether the increase of the area of the lymphatic vasculature at the cribriform plate is the result of lymphatic endothelial cell (LEC) proliferation of the newly formed lymphatic vessels at the cribriform plate, we collected single cell suspensions of cribriform plate areas for flow cytometry after 7 days of tMCAO. Among the singlets, we gated for cribriform plate LECs (cpLECs) which we defined as live, CD45^-^, CD31^+^, PDPN^+^ (Fig. 2A). We found that the number of cpLECs was significantly increased after tMCAO compared to sham groups (Fig. 2B). This data further corroborates our IHC analysis that lymphatics near the cribriform plate undergo lymphangiogenesis after tMCAO.

**Figure 2.**
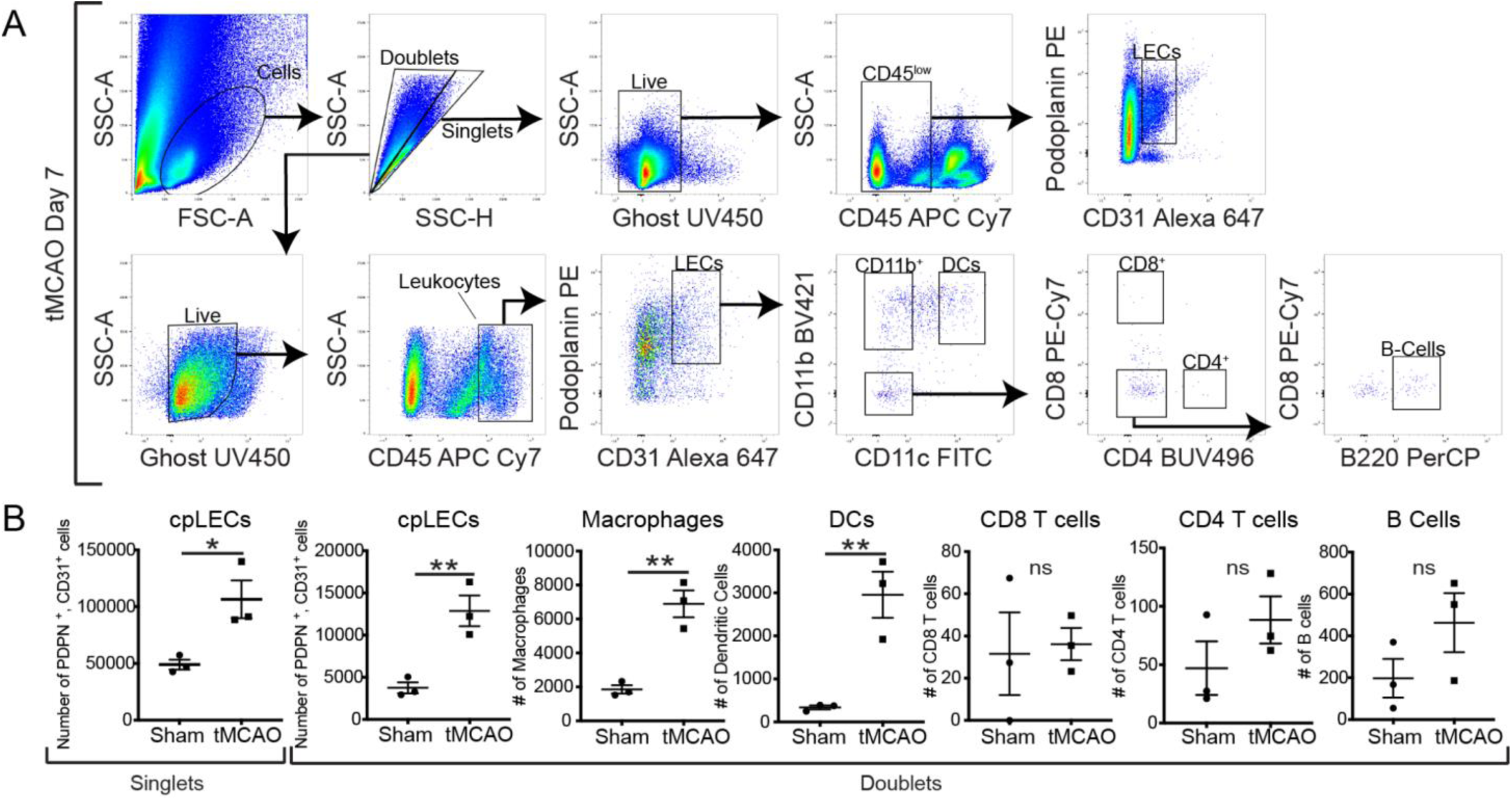
Lymphangiogenic vessels in the CP can interact with immune cells after tMCAO. **(A):** Cell-to-Cell interactions between cribriform plate lymphatic endothelial cells (cpLECs) and immune cells were studied by gating for live doublets from the cribriform plate cell suspensions. cpLECs were gated as doublets, live, and CD45^+^, CD31^+^, Podoplanin^+^. Macrophages were gated as CD45^hi^, CD11b^+^, CD11c^-^. Dendritic cells were gated as CD45^hi^, CD11b^+^, CD11c^+^. CD4 T cells were gated as CD45^hi^, CD11b^-^, CD11c^-^, CD4^+^. CD8 T cells were gated as CD45^hi^, CD11b-, CD11c-, CD8^+^. B cells were gated as CD45^hi^, CD11b^-^, CD11c^-^, CD4^-^, CD8^-^, B220^+^. cpLECs from singlets were gated as singlets, live, and CD45^-^, CD31^+^, Podoplanin^+^. **(B):** Quantitation of number of cpLECs in singlets, number of cpLECs in doublets, and number of immune cells in doublets near the cribriform plate between sham and tMCAO groups after 7 days (n=3 mice per group; mean ± SEM, *p < 0.05, **p ≤ 0.01, unpaired Student’s t-test).

To investigate if cpLECs interact with immune cells, we investigated cell to cell interaction by looking at doublets within the flow cytometry dataset (Giladi et al., 2020; Bendall, 2020). We analyzed the doublets between leukocytes and LECs near the cribriform plate during drainage after tMCAO. To accomplish this, we gated for LECs (CD31^+^, PDPN^+^) within CD45^+^ clusters and then further gated for macrophage (CD45^hi^, CD11b^+^, CD11c^-^), dendritic cells (CD45^hi^, CD11b^+^, CD11c^+^), CD8 T cells (CD45^hi^, CD11b^-^, CD11c^-^, CD8^+^), CD4 T cells (CD45^hi^, CD11b^-^, CD11c^-^, CD4^+^), and B cells (CD45^hi^, CD11b^-^, CD11c^-^, CD8^-^, CD4^-^, B220^+^) in both sham and tMCAO groups (Fig. 2A). We find that number of CD31^+^, PDPN^+^ cells, which we identified as cpLECs in doublets, was increased in tMCAO groups as well as macrophages and dendritic cells (Fig. 2B).

These data indicate that cpLECs interact directly with CD11b^+^, CD11c^-^ macrophages and CD11b^+^, CD11c^+^ dendritic cells. These interactions might contribute to the migration of macrophages and dendritic cells, as well as lymphangiogenesis near the cribriform plate after tMCAO.

In addition to drainage of immune cells, fluid drainage is another critical function of lymphatic vessels (Petrova and Koh, 2020). To test fluid drainage following tMCAO, we used T1-weighted MRI to confirm if these lymphangiogenic vessels can transport CSF after 7 days of tMCAO by administering gadolinium into the cisterna magna after taking baseline scans. On the dorsal view, we observed an increase in accumulation of gadolinium near the olfactory bulbs, which is directly adjacent to the cribriform plate (Fig. 3A). We also see an accumulation within the CLNs over time in the ventral view which correlates with lymphangiogenesis near the cribriform plate after 7 days of tMCAO (Fig. 3B). When we injected 10 μl of 10% Evans blue dye into the cisterna magna of tMCAO mice, the Evans blue dye in the coronal sections of cribriform plate regions co-localized with Lyve-1^+^ lymphatic vessels (Sup. Fig. 3A). Representative fluorescent intensity profile plot of Evans blue dye and Lyve-1 from a cross section of the merged image confirmed co-localization (dotted line) (Sup. Fig. 3B). Together this data suggests that CSF can be drained through lymphangiogenic vessels and reach CLNs after tMCAO.

**Figure 3.**
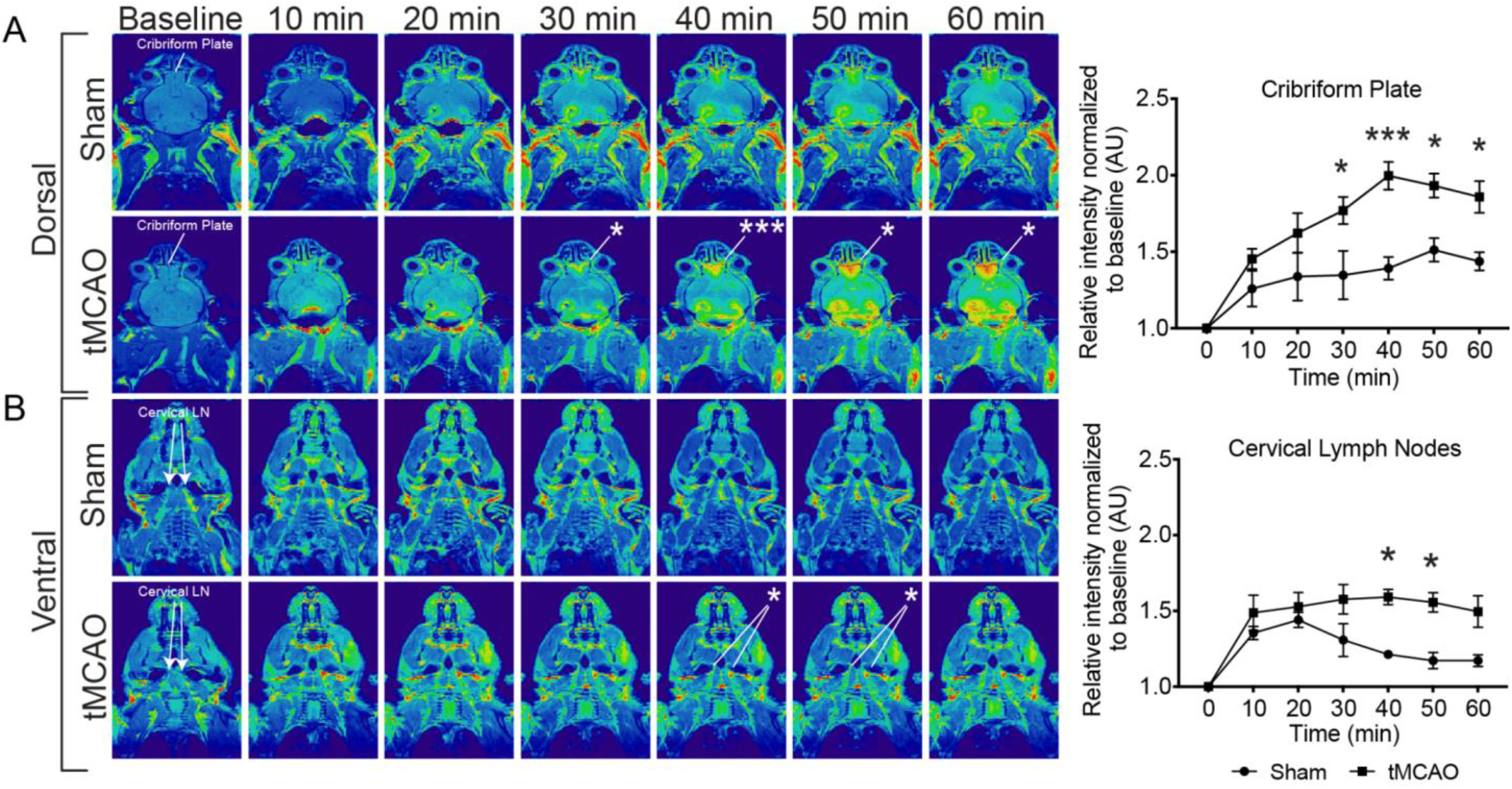
Fluid drains at the CP and CLNs after tMCAO. Representative T1-weighted MRI scan images of dorsal **(A)** and ventral **(B)** view of mice whole heads between sham and tMCAO. Baseline images were imaged before gadolinium injection and serial images over time were imaged after injection. Quantitation of average pixel intensity normalized to baseline of cribriform plate **(A)** and cervical lymph nodes **(B)** between sham and tMCAO mice (n=3 mice for sham, n=5 mice for tMCAO; mean ± SEM, *p ≤ 0.05, ***p ≤ 0.001, two-way ANOVA)

### Lymphangiogenesis occurs via the interaction of VEGF-C and VEGFR-3

The induction of lymphangiogenesis occurs via interaction of VEGF-C and VEGFR-3 expressed on LECs (Kajiya et al., 2009; Yan et al., 2017; Aspelund et al., 2014). After 7 days of tMCAO, cribriform plate areas of both sham and tMCAO groups were collected to measure VEGF-C protein expression during peak post-stroke lymphangiogenesis. The relative expression of VEGF-C by western blot was increased after tMCAO compared to sham with ß-actin as a loading control (Fig. 4A). Coronal sections of cribriform plate areas were stained using fluorescent-labeled antibodies of Lyve-1, CD11b, CD11c and VEGF-C. The images confirmed increased accumulation of immune cells and increased expression of VEGF-C near the cribriform plate after 7 days of tMCAO in mice (Fig. 4B). Orthogonal view of a cribriform plate section showed that Lyve-1^+^ lymphatics near the cribriform plate carry CD11b^+^, CD11c^+^ dendritic cells after tMCAO (Fig. 4C).

**Figure 4.**
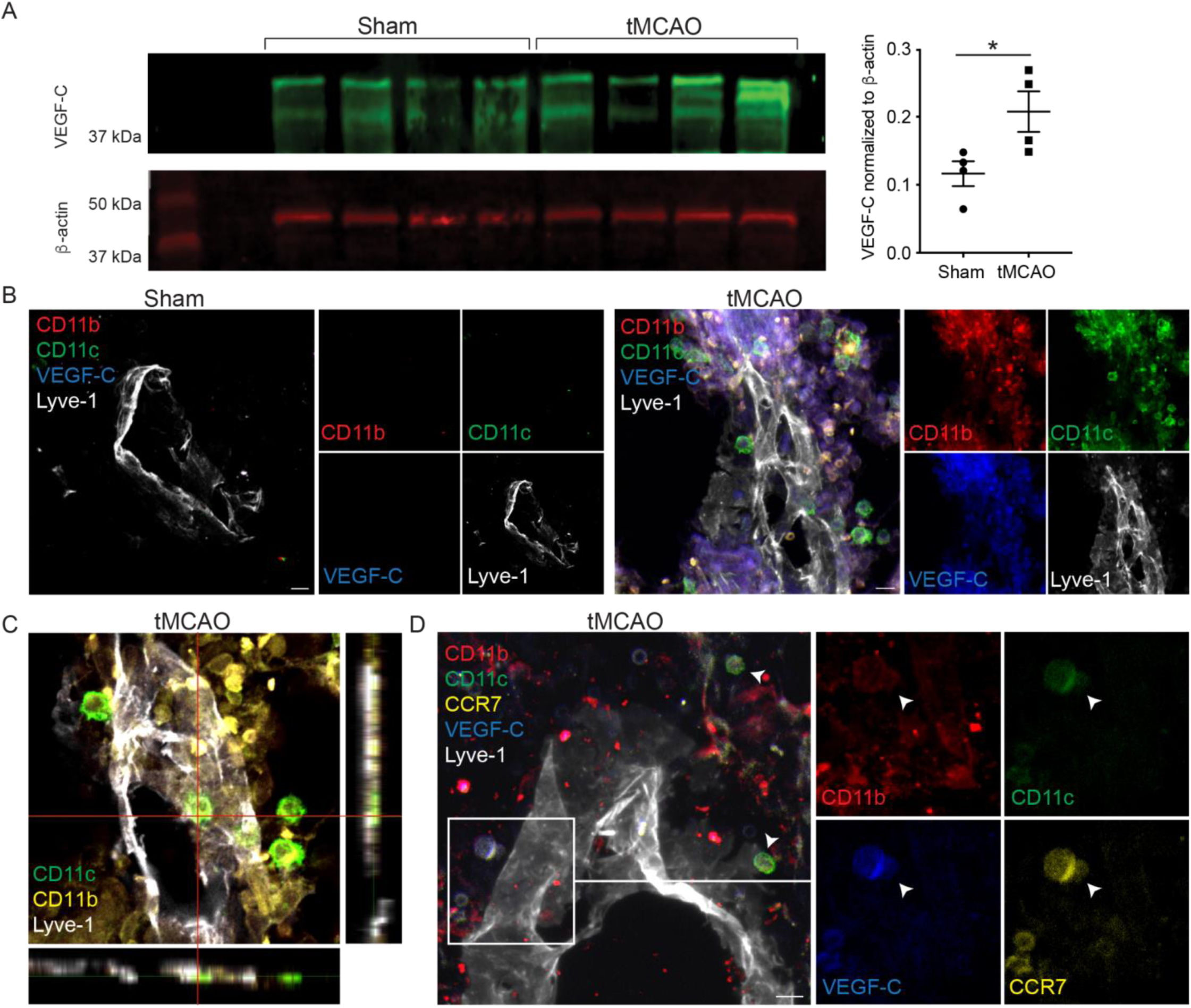
VEGF-C is upregulated at the CP after tMCAO. **(A):** VEGF-C expression was measured using Western Blot and β-actin was used as a loading control. Quantitation of relative VEGF-C amount normalized to β-actin (n=4 mice per group; mean ± SEM, *p < 0.05, unpaired Student’s t-test).**(B):** Representative sectional images of cribriform plate areas after immunolabeling with CD11b, CD11c, VEGF-C, and Lyve-1 to visualize immune cells and lymphatic vessels after 7 days of sham and tMCAO. Scale bars = 10 µm. **(C):** Representative orthogonal sectional view of cribriform plate after 7 days of tMCAO showing CD11b^+^, CD11c^+^ dendritic cells are bound to Lyve-1^+^ lymphatic vessels. Scale bar = 100 µm. **(D):** Representative sectional images of cribriform plate areas after immunolabeling with CD11b, CD11c, CCR7, VEGF-C, and Lyve-1 to visualize immune cells and lymphatic vessels after 7 days of tMCAO. Scale bars = 10 µm.

Additionally, we observed CD11b^high^, CD11c^+^, CCR7^+^ cells and CD11b^low^, CD11c^+^, CCR7^+^ cells co-localized with VEGF-C near the cribriform plate (Fig. 4D), showing that migratory dendritic cells are a potential source of VEGF-C at this site after 7 days of tMCAO.

### MAZ51 treatment improves motor recovery at earlier, but worsens at later time points after tMCAO

To further confirm the interaction between VEGF-C and VEGFR-3 after tMCAO, we used a VEGFR-3 inhibitor, MAZ51, to inhibit pro-lymphangiogenic signaling and to assess potential therapeutic effects (Kirkin et al., 2001, 2004). Either DMSO or MAZ51 was administered intraperitoneally on days 0, 2, 4, and 6 to control groups and experimental groups, respectively. Average Lyve-1^+^ vessel volumes near the cribriform plate were reduced after MAZ51 treatment in tMCAO groups, while MAZ51 did not induce further regression of lymphatics in sham groups (Fig. 5A). The COS and SSS of dural meningeal lymphatics showed decreased Lyve-1 expression after MAZ51 in tMCAO groups (Fig. 5B). Similarly, lymphatics in the CLNs showed regression of Lyve-1^+^ vessels after MAZ51 treatment (Fig. 5C). This data further confirms that lymphangiogenesis occurs via interaction of VEGF-C and VEGFR-3. Moreover, dilation of lymphatic vessels in the dural meningeal lymphatics may be induced by VEGF-C/VEGFR-3 interaction.

**Figure 5.**
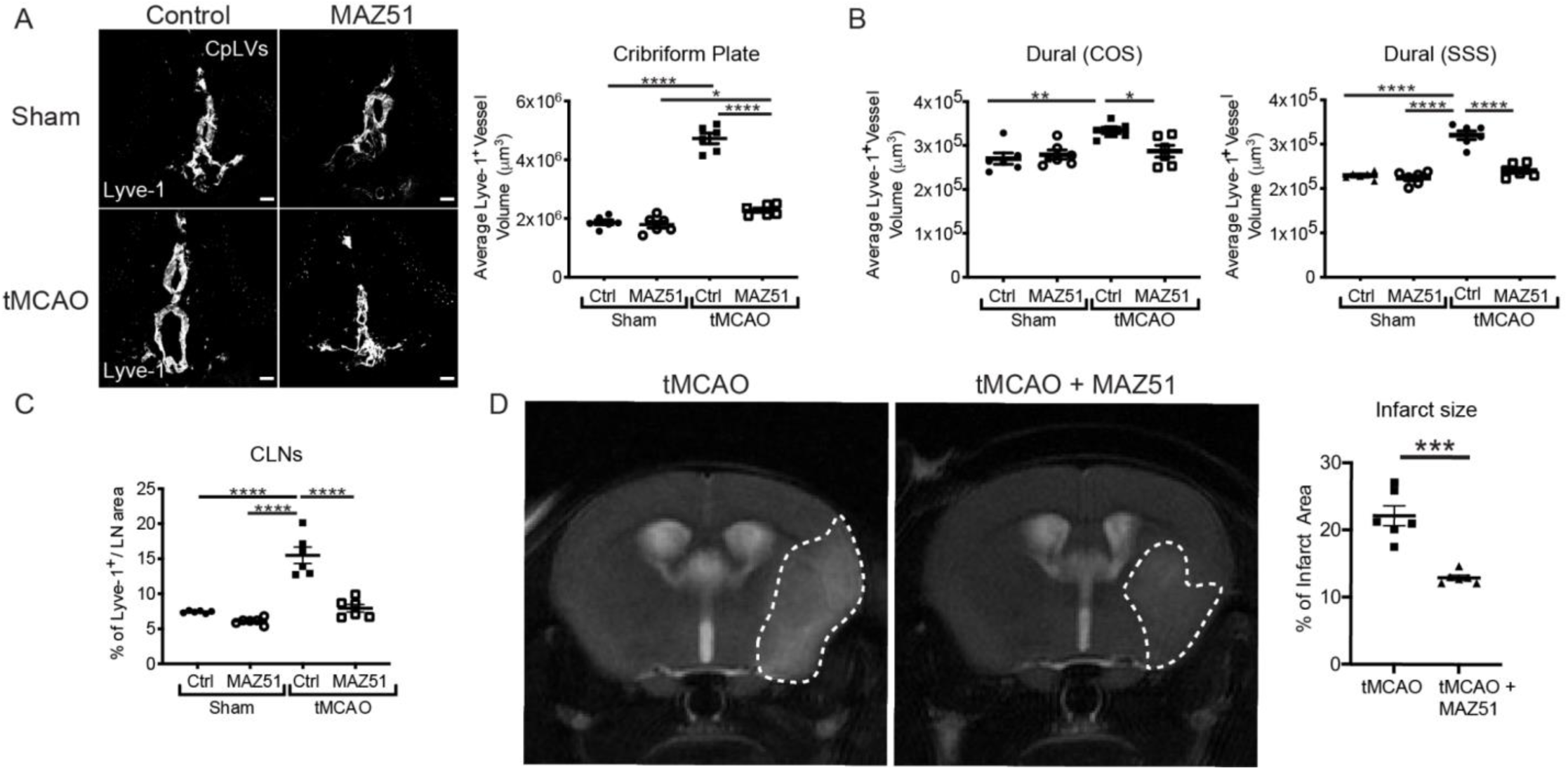
Post-stroke MAZ51 inhibits CP lymphangiogenesis and reduces brain infarct size. **(A):** Coronal sections of cribriform plate areas were stained with Lyve-1 fluorescent antibody after either control or MAZ51 treatment in Sham or tMCAO mice. Representative confocal images of lymphatic vessels near the cribriform plate. Scale bars = 100 µm for cribriform plate sections. Quantitation of average Lyve-1^+^ vessel volumes near the cribriform plate at Day 7 after stroke (n=6 mice per group; mean ± SEM, *p ≤ 0.05, ****p ≤ 0.0001, one-way ANOVA). **(B):** Quantitation of average Lyve-1^+^ vessel volumes near the COS and SSS of dural lymphatic vessels after either control or MAZ51 treatments between sham and tMCAO. (n=6 mice per group; mean ± SEM, *p ≤ 0.05, **p ≤ 0.01, ****p ≤ 0.0001, one-way ANOVA). **(C):** Quantitation of average Lyve-1^+^ vessel area within cervical lymph nodes. (n=6 mice per group; mean ± SEM, ****p ≤ 0.0001, one-way ANOVA). **(D):** T2-weighted MRI scan images of brain sections between tMCAO with control or MAZ51 treatment at day 7. Representative images of MRI brain scans. White dashed lines mark brain infarcts which are quantified into a graph (n=6 mice per group; mean ± SEM, ***p ≤ 0.001, unpaired Student’s t-test)

In order to test the role of CP lymphangiogenesis in the pathological outcomes following stroke, we performed T2-weighted MRI imaging to measure brain infarction following i.p. administration of MAZ51. Interestingly, seven days following tMCAO, MAZ51-treated mice showed decreased infarction areas compared to control-treated tMCAO mice (Fig. 5D). During MAZ51 treatments, three behavior tests such as ladder rung test, open-field, and rotarod, were also conducted on days 1, 3, 5, and 7 after sham or tMCAO (Sup. Fig. 4A).

On the ladder rung test, MAZ51-treated tMCAO mice showed improvements by making less foot faults starting from day 3 compared to control-treated tMCAO mice (Sup. Fig. 4B). In the open-field tests, each mouse was placed in the center of the box and their movements were tracked. Quantitation of the duration in the center of the box showed there was a trend indicating that MAZ51-treated tMCAO mice stayed less time in the center compared to control-treated tMCAO mice on day 1, although this was not statistically significant (Sup. Fig. 4C-D). Broadly, failure to exit the center zone after initial placement was associated with a high level of motor impairment in stroke mice during the open field test, as mice were unable to travel in a straight line toward the peripheral wall (Sup. Fig. 4C; Trackmap). In the rotarod test, control-treated tMCAO mice showed improvements at day 7, while MAZ51-treated tMCAO mice did not (Sup. Fig. 4E). In summary, MAZ51-treated tMCAO mice showed better locomotor ability as ladder rung tests and open-field tests as well reduced brain infarct areas, starting from as early as day 1. However, these MAZ51-treated tMCAO mice showed hindered natural recovery along with worse coordination and balance at day 7 in rotarod tests. This result may correlate with regressed lymphangiogenesis near the cribriform plate and lymphatic dilation of dural meningeal lymphatics after MAZ51 treatment (Sup. Fig. 4F).

### Delivering VEGF-C after tMCAO does not induce changes in lymphatics but worsens brain infarct

VEGF-C administration has been proposed to diminish brain edema, increase brain drainage and promote regeneration following neuroinflammatory damage to the brain (Tsai et al., 2022; Matta et al., 2021). To test whether this treatment would be beneficial following ischemic damage to the brain, we induced 60-minute ischemia followed by reperfusion using the tMCAO model. We then administered VEGF-C156S or PBS through the common carotid artery using a microcatheter on day 0 of the tMCAO (Fig. 6A). VEGF-C156S specifically activates VEGFR-3, unlike endogenous forms of VEGF-C which can also bind to VEGFR-2 (Joukov et al., 1998). Quantitation of Lyve-1^+^ vessels showed there was no further increase in lymphatics near the cribriform plate, COS and SSS of dural meningeal lymphatics, or in CLNs after VEGF-C156S treatment (Fig. 6B, Sup. Fig. 5A-C). However, brain infarction was significantly increased after VEGF-C156S treatment at both day 3 and 7 compared to control-treated tMCAO mice (Fig. 5C). To better understand the elevation of infarct size, we investigated alterations in peripheral immune cell populations as they may have been influenced from VEGF-C15S administration. Interestingly, we found that around the subfornical vessels of the brain, a commonly reported site of immune cell infiltration into the CNS (Schulz and Engelhardt, 2005; D’Mello et al., 2009), there was an increased number of CD45^+^ leukocytes in the perivascular regions of the vessels (Fig. 5D). Together this data suggests that post-stroke VEGF-C delivery could exacerbate stroke damage, elevate neuroinflammation, and promote immune cell aggregation at barrier sites in the brain while not directly affecting lymphatics surrounding the CNS.

**Figure 6.**
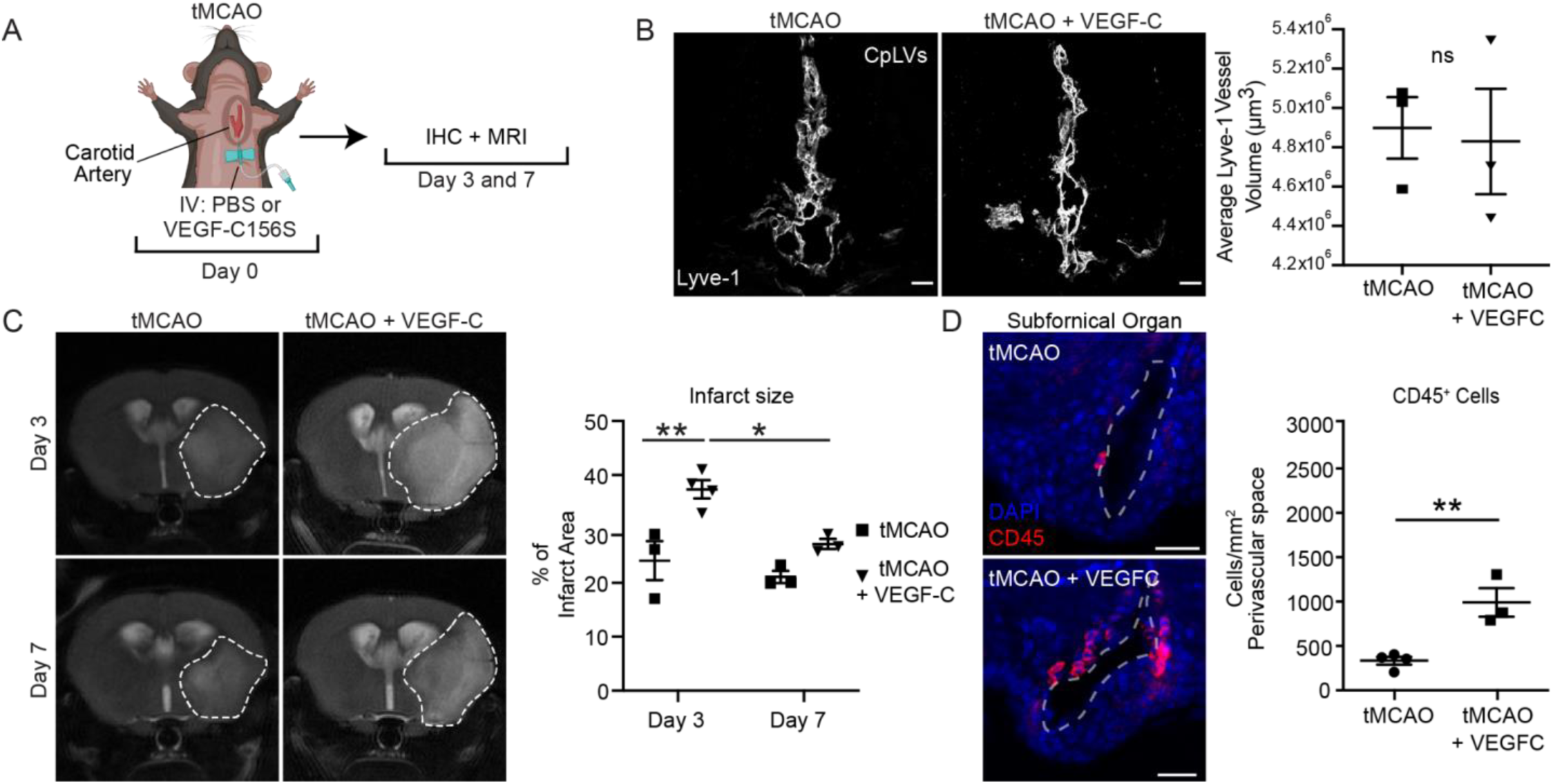
Injecting VEGF-C after tMCAO does not induce additional changes in lymphatics, but worsens the brain infarct. **(A):** Experimental design for administering control or VEGF-C156S through artery after reperfusion of tMCAO (day 0) and euthanization at day 3 and 7 for decalcification steps. **(B):** Coronal sections of cribriform plate areas were stained with Lyve-1 fluorescent antibody after either control or VEGF-C156S treatment of tMCAO mice at day 7. Representative confocal images of lymphatic vessels near the cribriform plate. Scale bars = 100 µm for cribriform plate sections. Quantitation of each image (n=3 mice per group; mean ± SEM, unpaired Student’s t-test). **(C):** T2-weighted MRI scanned images of brain sections between tMCAO and tMCAO with VEGF-C156S treatment at day 3 and 7. Representative images of MRI brain scans. White dashed lines mark brain infarcts which are quantified into a graph (n=3 mice per group, n=4 mice for tMCAO with VEGF-C156S at day 3; mean ± SEM, *p ≤ 0.05, **p ≤ 0.0, two-way ANOVA with mixed effects). **(D):** Coronal brain sections were stained with CD45 fluorescent antibody (Alexa 647) at Day 3 post stroke/VEGF-C treatment. Subfornical vessels near the third ventricle were analyzed for perivascular leukocyte cell number. (n=4 mice for tMCAO, n=3 mice for tMCAO with VEGF-C156S; mean ± SEM, **p ≤ 0.01, unpaired Student’s t-test). Scale bars = 20 µm

## Discussion

Here we showed for the first time that lymphatic vessels near the cribriform plate undergo lymphangiogenesis after tMCAO and that these newly formed vessels can drain CSF as well as retain immune cells during stroke. As ischemic areas undergo necrosis and apoptosis due to deprivation of oxygen and nutrients, pro-inflammatory cytokines and chemokines are released which recruit immune cells to the injury site (Gesuete et al., 2016). Simultaneously, cerebral edema can occur as one of the complications of ischemic stroke (Dostovic et al., 2016). Since the brain is enclosed within the skull, increased intracranial pressure exerted by the brain and CSF can lead to secondary brain ischemia through the reduction of cerebral blood flow and increasing tissue hypoxia (Jeon et al., 2014). This can ultimately cause death in some stroke patients (Jeon et al., 2014; Thorén et al., 2017). Effective drainage of fluid from the CNS especially during inflammation would be necessary in order to prevent the worst outcomes. Multiple routes that drain fluid and cells in the CNS have been re-characterized recently (Engelhardt et al., 2017; Proulx, 2021; Laaker et al., 2023). However there are still controversial debates regarding each route’s contribution to drainage of CSF, antigen, and immune cells during both steady state and neuroinflammation. Several groups have shown that lymphatics near the cribriform plate play significant roles in drainage of CSF and immune cells to CLNs to coordinate immune responses (Hsu et al., 2019; Walter et al., 2006; Pashenkov et al., 2003; Cserr et al., 1992; Mollanji et al., 2001). Additionally, recent characterization of the cribriform plate lymphatics in mice have revealed discontinuity of the E-cadherin+ arachnoid layer at this region, potentially allowing cribriform lymphatic vessels more permissible access to fluid, antigen, and cells draining in the subarachnoid space around olfactory nerves (Spera et al., 2023; Hsu et al., 2022).

The increased numbers of loops and sprouts at the cribriform plate correlated with increased Lyve-1^+^ vessel volumes, indicating that lymphangiogenesis near the cribriform plate occurs at around day 3 and appears to regress by day 14. Flow cytometry data further showed that there was an expansion of the number of LECs after tMCAO. However, dural meningeal lymphatic appears to undergo vessel dilation after tMCAO. Dural meningeal lymphatics have shown an increase of diameters in response to VEGF-C which is naturally elevated in CSF after stroke due to increased interstitial fluid (ISF) pressure and lymphedema (Da Mesquita et al., 2018; Esposito et al., 2019; Gu et al., 2001; Rutkowski et al., 2006). Increased aquaporin 4 polarization on astrocytes after stroke can further contribute to cerebral edema (Zador et al., 2009). Other groups have shown that impairment of dural meningeal lymphatics affected clearance of macromolecules, but not ISF pressure which indicates there is another route contributing to removal of extra fluid in the brain during inflammation (Da Mesquita et al., 2018; Aspelund et al., 2015). Furthermore, we found that only the cervical lymph nodes experienced post-stroke lymphangiogenesis, implying the greater connection to CNS fluid and cell drainage pathways may be necessary for driving post-stroke lymphatic vessel growth in the periphery.

The interaction of VEGF-C and VEGFR-3 has been indicated as a key driver of lymphangiogenesis (Baluk et al., 2005; Jussila and Alitalo, 2002). We showed an upregulation of VEGF-C protein expression near the cribriform plate after stroke induction, and additionally we identified that VEGF-C was produced by CD11b^+^ CD11c^+^ dendritic cells. Since some of the dendritic cells also expressed CCR7, populations of migratory dendritic cells may produce VEGF-C during lymphangiogenesis near the cribriform plate after tMCAO. When lymphangiogenesis was inhibited using MAZ51, MAZ51 did not affect the sham groups which indicate that there may be a baseline threshold in lymphatic density. Additionally, MAZ51 may inhibit recruitment and activation of immune cells. Esposito et al. observed LEC proliferation and activation of macrophages in the CLNs within 24 hours after MCAO in rats. MAZ51 treatment reduced pro-inflammatory macrophages and LEC activation which also resulted in smaller brain infarcts. They further showed that VEGF-C/VEGFR-3 interaction increased inflammatory responses in LECs in co-cultured macrophages in *in vitro* experiments (Esposito et al., 2019). Combined with our behavioral data showing improved locomotor ability at earlier time points, inhibiting lymphangiogenesis with MAZ51 to reduce recruitment of activated pro-inflammatory immune cells to the brain may be beneficial at an earlier period after tMCAO.

However, Breslin et al. showed MAZ51 treatment decreased lymphatic phasic activities and function of lymphatic pumps (Breslin et al., 2007). In our study, MAZ51-treated tMCAO mice failed to show natural recovery in rotarod tests after 7 days. Interestingly, this is when we noticed a peak of lymphangiogenesis near the cribriform plate after tMCAO. As both lymphangiogenesis and lymphatic dilation were inhibited with MAZ51, the cerebral edema may further suppress other areas of the brain and induce secondary damages (Jeon et al., 2014; Wijdicks et al., 2014). Thus, prolonged inhibition of lymphangiogenesis or dilation may affect fluid drainage and recovery after tMCAO, but further analysis is needed to dissect the exact mechanism.

Conversely, administering VEGF-C156S directly to the brain through the common carotid artery resulted in larger brain infarcts. VEGF-C is a chemokine regulated by infiltrating immune cells and pro-inflammatory cytokines such as IL-1ß and TNF-α, unlike VEGF-A that is regulated by hypoxia (Jussila and Alitalo, 2002; Enholm et al., 1997; Ristimäki et al., 1998). It is involved in not only lymphangiogenesis but also recruiting immune cells especially those that express VEGFR-3 such as macrophages and dendritic cells (Li et al., 2016). Thus, it is possible that VEGF-C156S resulted in increased immune cells recruitment toward core regions of brain infarcts after tMCAO which caused bigger infarcts compared to control-treated tMCAO mice. These results contrast recent reports that suggest that VEGF-C delivery after stroke may offer recovery benefits (Matta et al., 2021; Tsai et al., 2022). One explanation for this divergence in effects could be the result of differences in stroke model. Tsai et al., indicated therapeutic enhancement of meningeal lymphatics via VEGF-C156S allowed for greater clearance of hematomas, in model intracerebral hemorrhage not tMCAO (Tsai et al., 2022). As a result, beneficial manipulation of post-stroke VEGF-C / VEGFR-3 signaling could be highly context dependent, unique to stroke type or even stroke location. Additionally, as our data suggests, there could be a specific therapeutic window in which VEGF-C/VEGFR-3 modifying therapy should be given, much like current tPA treatment protocols. In agreement with this, pretreatment with VEGF-C was shown to yield improved stroke outcomes (Simoes Braga Boisserand et al., 2023)

In summary, this paper showed that lymphatics near the cribriform plate undergo lymphangiogenesis, while dural meningeal lymphatics are dilated after tMCAO in mice. The lymphangiogenesis was driven by the interaction of VEGFR-3 and VEGF-C which is produced by migratory dendritic cells. The VEGFR-3 inhibitor blocked lymphangiogenesis after tMCAO and created smaller brain infarcts. Based on the behavioral test results, inhibition of VEGFR-3-dependent lymphangiogenesis after tMCAO seems beneficial at earlier time points, but may have long-term consequences on recovery depending on length of treatment. On the other hand, administering VEGF-C into the brain directly after stroke induces bigger brain infarct areas. More research is needed to investigate more optimal timings of potential VEGFR-3 and VEGF-C therapeutic manipulations, as well as more specific and safe targeting of meningeal lymphatic vessels. These data emphasize caution when targeting the VEGF-C/VEGFR-3 pathway in promoting post-stroke recovery in spite of the impressive effects of VEGF-C/VEGFR-3 mediated pathway on tissue regeneration following stroke.

## Materials and Methods

### Animals

Male C57BL/6J wild type was purchased from Jackson Laboratories and housed in University of Wisconsin at Madison Breeding Core and Research Services. Animals were kept in a pathogen-free facility with 12 hours of each dark and light cycle and access to food and water. All experiments were conducted in accordance with guidelines from National Institutes of Health and the University of Wisconsin-Madison Institutional Animal Care and Use Committee.

### tMCAO

10-12 weeks old male mice (25-29 g) were used for all transient middle cerebral artery occlusion (tMCAO) surgery. In agreement with the STAIR criteria (Liu et al., 2009), core temperature was maintained between 36-37 °C during surgery with a heating pad and post-surgery mice recovered in a temperature-controlled chamber. Mice were anesthetized using isoflurane while providing oxygen during occlusion and reperfusion surgery. During occlusion, midline incision was made and the common carotid artery (CCA) on the right side was isolated from the vagus nerve. External carotid artery (ECA) and internal carotid artery (ICA) were identified from the bifurcation of CCA. Both bottom of the CCA and ECA were permanently ligated. A temporary knot was made near the bifurcation of CCA. A microvascular clip was made on CCA and a 6.0 nylon monofilament (3021910; Doccol Corp) was inserted into the middle cerebral artery (MCA) while loosening the temporary knot carefully. Temporary knot was tightened again, and the occlusion was initiated by advancing the filament ∼ 9 to 9.5 mm to block MCA blood flow for 60 minutes. After 60 minutes, mice were anesthetized, and the filament was removed. The temporary knot was tightened as a permanent ligation. For sham mice, all surgical operations were the same, but the filament was not inserted. According to our power calculations, n = 3–10 sex- and age-matched mice were used for each with group assignments randomized.

### Flow Cytometry

After 7 days of tMCAO, single cell suspensions from cribriform plate were resuspended in FACS buffer (pH 7.4, 0.1 M PBS, 1 mM EDTA, 1% BSA) and immunolabeled with the appropriate conjugated antibodies for 30 minutes at 4°C. The stained cells were washed with FACS buffer and fixed in 4% PFA (4% paraformaldehyde, 0.1 M PBS). Data was acquired using BD LSR II Flow Cytometer (BD Biosciences) and analyzed using FlowJo software.

### Histology

Mice were anesthetized with isoflurane and transcardially perfused with 1X PBS followed by 4% PFA. Then, mice were decapitated and the whole heads without skin were fixed in 4% PFA overnight at 4°C. Then the whole heads were transferred to 14% EDTA for 7 days to decalcify followed by cryoprotection step in 40% sucrose for 7 days at 4°C. Then, the heads were embedded in Tissue-Tek OCT Compound, frozen in dry ice, and stored at -80°C. Leica CM1800 (Leica Biosystems) cryostat was used to section each head into 60 μm tissue slides for cribriform plate area and 30 μm for brains and lymph nodes. Each section was mounted on Superfrost Plus microscope slides and stored at -80°C till staining.

### Infarct Area Determination

Brain infarction was screened 4.7 T small animal MRI (Agilent Technologies Inc., Santa Clara, CA) and acquired with VnmrJ (Agilent Technologies) on day 3, 7, and 14. Animals were anesthetized using isoflurane through a nose-cone during imaging. T2-weighted MRI scans were measured under the following parameters: TR= 3500 ms, thickness= 1.0 mm, resolution= 192×192, averages = 11. Each image was analyzed using FIJI software. Ischemic infarct area in percentage was calculated by dividing infarction area with total area of each brain section.

### Drainage Study

10 μl of 10% Evans Blue dye was injected into the cisterna magna using a Hamilton syringe at a rate of 2 µl/minutes. After allowing the dye to circulate for 30 minutes, the mouse was euthanized and the whole head was analyzed for dye distribution around the cribriform plate. MRI was done with a 4.7 T small animal MRI (Agilent Technologies Inc., Santa Clara, CA) and acquired using VnmrJ (Agilent Technologies). 2D T1-weighted MRI scans were used to detect gadolinium under the following parameters: TR= 688 ms, TE= 11.26 ms, thickness= 0.5 mm, resolution= 128×128, averages=7. These resulted in a time scan of about 5 min. We repeated this scan post injection for 1 hour. Animals were anesthetized using isoflurane through a nose-cone and 10 ul of gadolinium was injected into the cisterna magna at a rate of 2 μl/min using a Hamilton syringe. Respiratory rates were monitored throughout the scans. A baseline scan was acquired prior to gadolinium injection with the same settings. Images were processed and analyzed using FIJI software.

### Immunohistochemistry and Confocal Microscopy

For immunohistochemistry, sections were rehydrated with 1X PBS for 10 minutes, and blocking solution (1% BSA and 0.1% Triton-X in 1X PBS) for 60 minutes. Sections were then incubated with the appropriate primary antibodies in blocking solution at 4°C overnight in a humidified chamber. Sections were washed 3 times with 1X PBS for 10 minutes each. As necessary, sections were incubated with the appropriate secondary antibodies at room temperature for 120 minutes and washed afterward. Then, each section was mounted with Prolong Gold mounting medium with DAPI and images were acquired using an Olympus Fluoview FV1200 confocal microscope with 4x, 10x, or 20x objectives. Each image was analyzed using FIJI software (version 2.3.0/1.53q). Detailed list of reagents and antibodies are provided in Supplementary Table 1.

### Quantification of loops, sprouts, and lymphatic vessel diameter

Lymphatic vessel loops and sprouts were counted under identical confocal settings and FIJI software. Lymphatic vessel diameter was assessed with 50 measurements per meningeal sample in each group along either COS or SSS and then averaged together. All measurements were done by an independent blinded experimenter.

### Western Blot

Mice were anesthetized with isoflurane and transcardially perfused with 1X PBS. Harvested brains were doused in RIPA buffer (pH 7.5, 25 mM Tris-Cl, 150 mM NaCl, 1mM EDTA, 1% Triton-X 100, 0.1% SDS) and the appropriate dilution of protease and phosphatase inhibitors. The grinded tissues were sonicated and stored at -80°C. Protein levels were assessed with the Li-Cor Odyssey CLx infrared imaging system. Detailed list of reagents and antibodies are provided in Supplementary Table 1.

### Drug Administration

MAZ51 was dissolved in DMSO and intraperitoneally injected at 10 mg/kg of mouse weight on day 0 (after reperfusion of tMCAO), 2, 4, 6. Control mice received equivalent volumes of DMSO intraperitoneally on those same days. VEGF-C156S was dissolved in 1X PBS and 3 ug was administered through a micro-catheter inserted in CCA on day 0 after reperfusion of tMCAO. Control mice received equivalent volumes of 1X PBS using the same method on day 0.

### Behavior Tests

Post-ischemic motor functions were measured using rotarod (LE8205; Panlab Harvard Apparatus), ladder rung test (LE780; Panlab Harvard Apparatus), and open-field (LE802S from Panlab Harvard Apparatus). Rotarod tests were conducted for 3 minutes on a rotating cylinder with constant speed at 8 rpm.

Ladder rung test was conducted until each mouse crossed the tapered 100 cm beam and number of foot faults were counted. Open-field test was used to observe spontaneous locomotor functions in a 45 (W) x 45 (D) x 40 (H) cm box for 5 minutes. Open-field test was analyzed using Smart 3.0 software (Panlab Harvard Apparatus).

All mice were trained with three tests 2 consecutive days before the surgery and randomized for sham or tMCAO surgery on day 0. Tests were measured on days 1, 3, 5, and 7 after MAZ51 treatment. All behavioral equipment was wiped with 70% ethanol between mice.

### Statistical Analysis

Statistical analysis was performed with GraphPad Prism 6.0 Software. When results of two groups were compared, unpaired Student’s t-test was used. When results of three or more groups were compared, one-way ANOVA was used. When results were compared across different time points, two-way ANOVA using Sidak’s multiple comparisons was used. For all statistical tests, the data are portrayed as mean ± standard error of the mean (SEM) and the significance is portrayed as: NS (p>0.05), *(p<0.05), **(p<0.01), ***(p<0.001), ****(p<0.0001).

## Acknowledgments

We thank Khen Macvilay for his expertise in flow cytometry, Laura Schmitt-Brunold for her expertise in molecular biology, and all our laboratory members for insightful comments on this work. We would like to thank the UW Small Animal Imaging Facility (SAIF) supported by the UWCCC grant P30CA014520 to use its facilities and services. This work was supported by the National Institutes of Health grants NS10847 and NS103506 awarded to ZF, HL128778 awarded to MS, the Neuroscience Training Program T32-GM007507 awarded to MH and CL, and AHA grant 915125 awarded to CL.

## Contributions

Y.H.C, M.S., and Z.F. conceptualized the experiments and reviewed, revised the manuscript. Y.H.C, performed the experiments, generated the figures, analyzed the data, and wrote the manuscript. M.Hsu assisted with the Evans Blue experiments, IHC analysis, and assisted with the FACS. C.L. assisted with behavioral experiments, IHC analysis, and helped revise/format the manuscript. M.H. assisted with the FACS. H.Y. assisted with IHC analyses. P.C., L.J., B.S., and K.W. assisted with behavioral analysis. M.S.and Z.F. are co-senior authors. All authors reviewed the paper.

## Competing Interests

The authors declare no competing interests.

---

**Supplementary Table 1.** Antibody List.

| Antibody | Conjugate | Clone | Source | Catalog ID | RRID |
| --- | --- | --- | --- | --- | --- |
| CD31 | Alexa 647 | 390 | BD | 563608 | AB_2738313 |
| CD45 | APC Cy7 | 30-F11 | Biolegend | 103115 | AB_312980 |
| Podoplanin | PE | eBio8.1.1 | Thermo Fisher Scientific | 12-5381-80 | AB_1907440 |
| CD11c | FITC | N418 | Biolegend | 117305 | AB_313774 |
| CD11b | BV421 | M1/70 | BD | 562605 | AB_11152949 |
| CD4 | BUV 496 | GK1.5 | BD | 612952 | AB_2813886 |
| B220 | PerCP | RA3-6B2 | BD | 553093 | AB_394622 |
| CD8 | PE-Cy7 | 53-6.7 | Biolegend | 100721 | AB_312760 |
| Lyve-1 | Alexa 660 | ALY7 | Thermo Fisher Scientific | 50-0443-80 | AB_10598060 |
| Lyve-1 | Alexa 488 | ALY7 | Thermo Fisher Scientific | 53-0443-82 | AB_1633415 |
| CD197 (CCR7) | PE | 4B12 | Thermo Fisher Scientific | 12-1971-82 | AB_465905 |
| VEGF-C (Unconjugated - Rabbit) | N/A | N/A | Thermo Fisher Scientific | PA5-29772 | AB_2547246 |
| Goat Anti-Rabbit | Alexa 405 | N/A | Thermo Fisher Scientific | A-31556 | AB_221605 |
| CD45 | Alexa 647 | 104 | Biolegend | 109818 | AB_492870 |
| CD11b | PE | M1/70 | Thermo Fisher Scientific | 12-0112-82 | AB_2734869 |
| CD11c | Alexa 488 | N418 | Thermo Fisher Scientific | 53-0114-80 | AB_469903 |
| CD11b | APC-eFlour 780 | M1/70 | eBioscience | 47-0112-82 | AB_1603193 |
| CD11b | APC/Cy7 | M1/70 | BioLegend | 101226 | AB_830642 |
| Mouse β-actin | N/A | mAbcam8226 | Abcam | ab8226 | AB_306371 |
| IRDye Donkey Anti-Rabbit IgG | 800CW | N/A | Li-Cor | 926-32213 | AB_621848 |
| IRDye Donkey Anti-Mouse IgG | 680RD | N/A | Li-Cor | 926-68072 | AB_10953628 |
| VEGF-C (Unconjugated - Rabbit) | N/A | N/A | Abcam | ab9546 | AB_2241408 |

**Supplementary Table 2.** Reagent List.

| Reagents | Source | Catalog ID |
| --- | --- | --- |
| Recombinant Human VEGF-C(Cys156Ser), CF | R&D Systems | 752-VC-025/CF |
| VEGFR-3 Kinase Inhibitor, MAZ51 | EMD Millipore | 676492 |
| Evans Blue | Sigma Aldrich | E2129-10G |
| Gadolinium | Bracco Diagnostics | NDC 0270-5164-14 |
| Protease and Phosphatase Inhibitor Cocktail | Thermo Fisher Scientific | 78440 |
| HBSS | Corning | 21-021-CV |
| RPMI-1640 | Corning | 10-040-CV |
| Dimethyl Sulfoxide | Thermo Fisher Scientific | D128-500 |
| Collagenase/Dispase | Sigma-Aldrich | 1026938001 |
| Ethylenediaminetetraacetic Acid | Thermo Fisher Scientific | BP118-500 |
| ProLong Gold Antifade Mountant with DAPI | Thermo Fisher Scientific | P36935 |
| ProLong Gold Antifade Mountant | Thermo Fisher Scientific | P36934 |
| UltraComp eBeads Compensation Beads | Thermo Fisher Scientific | 01-2222-42 |
| PI-132 microcatheter 150mm | Doccol Corporation | PI-SC1-132-097-150 |
| Sucrose, ACS Grade | Grainger | 31GD80 |
| Paraformaldehyde $\geq 96\%$, extra pure | VWR | 103528-632 |
| Ghost (UV540) | Tonbo | 13-0879-T500 |

## Notes

### Competing Interest Statement

The authors have declared no competing interest.

