## Supplementary_Figures for "Dual role of Vascular Endothelial Growth Factor-C (VEGF-C) in post-stroke recovery"

**Supplementary Figure 1. Number of lymphatic loops and sprouts near the cribriform plate increases after tMCAO.**

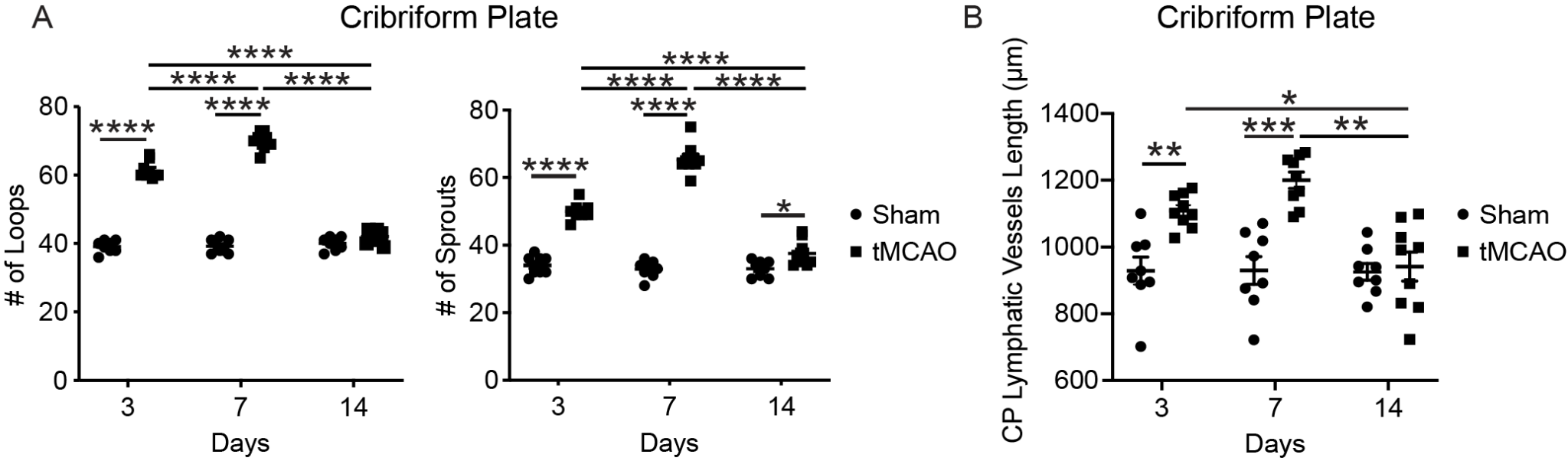

**Supplementary Figure 1. Number of lymphatic loops and sprouts near the cribriform plate increases after tMCAO.**

**(A):** Quantitation of number of loops and sprouts near the cribriform plate after sham and tMCAO at day 3, 7, and 14 (n=8 mice for sham, n=9 mice for tMCAO; mean ± SEM, *p ≤ 0.05, **p ≤ 0.01, ***p ≤ 0.001, ****p ≤ 0.0001, two-way ANOVA).

**(B):** Quantitation of length of lymphatics near the cribriform plate after sham and tMCAO at day 3, 7, and 14 (n=8 mice for sham, n=9 mice for tMCAO; mean ± SEM, *p ≤ 0.05, **p ≤ 0.01, ***p ≤ 0.001, ****p ≤ 0.0001, two-way ANOVA).

**Supplementary Figure 2. Lymphatic dilation occurs in dural meningeal lymphatics after tMCAO.**

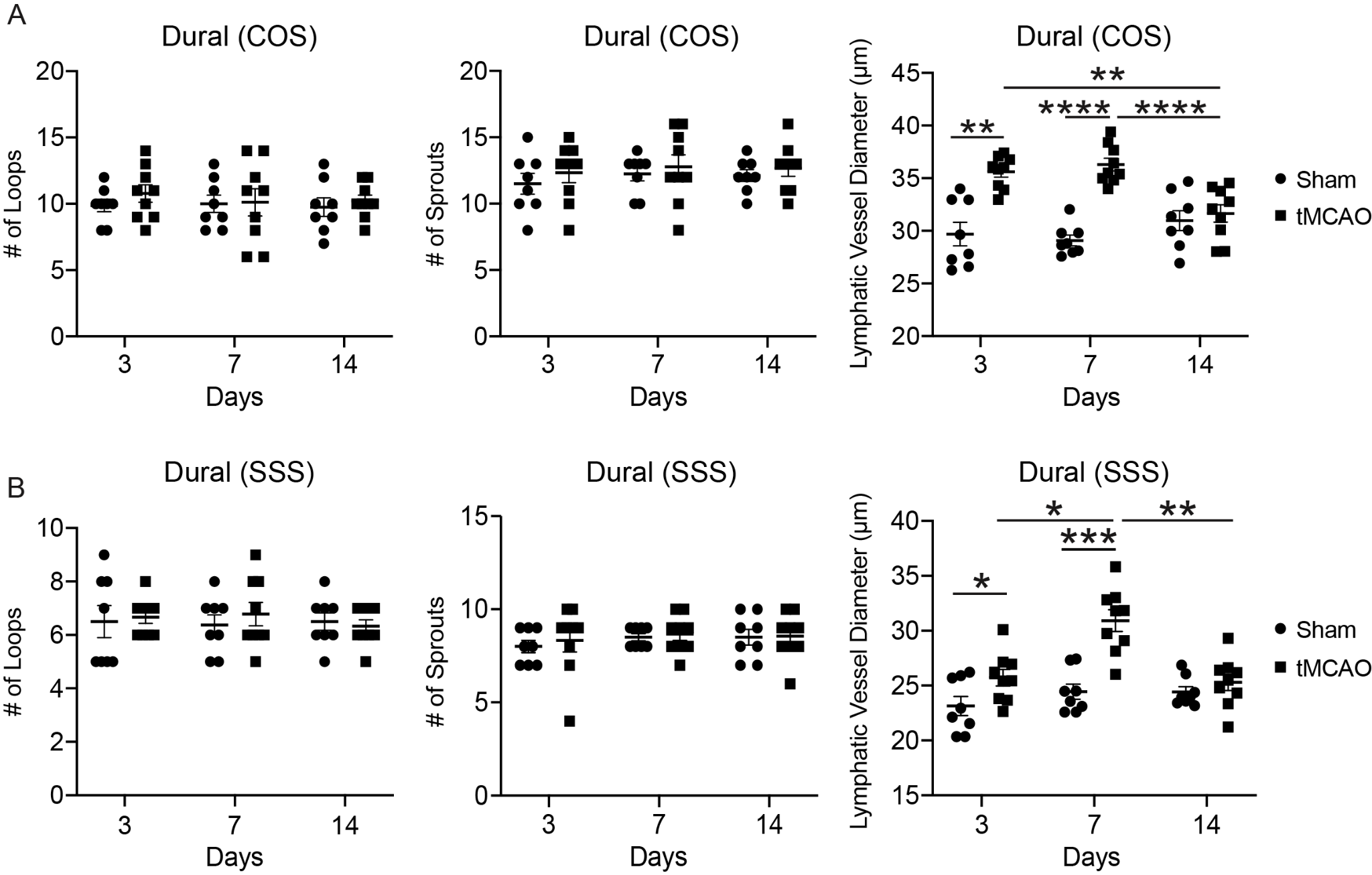

**Supplementary Figure 2. Lymphatic dilation occurs in dural meningeal lymphatics after tMCAO.**

**(A):** Quantitation of number of loops, number of sprouts, and lymphatic vessel diameters of confluence of sinuses (COS) of dural lymphatic vessels after sham and tMCAO at day 3, 7, and 14 (n=8 mice for sham, n=9 mice for tMCAO; mean ± SEM, *p ≤ 0.05, **p ≤ 0.01, ***p ≤ 0.001, ****p ≤ 0.0001, two-way ANOVA).

**(B):** Quantitation of number of loops, number of sprouts, and lymphatic vessel diameters of superior sagittal sinus (SSS) of dural lymphatic vessels after sham and tMCAO at day 3, 7, and 14 (n=8 mice for sham, n=9 mice for tMCAO; mean ± SEM, *p ≤ 0.05, **p ≤ 0.01, ***p ≤ 0.001, ****p ≤ 0.0001, two-way ANOVA).

**Supplementary Figure 3. Lymphangiogenic vessels in the cribriform plate can drain CSF after tMCAO.**

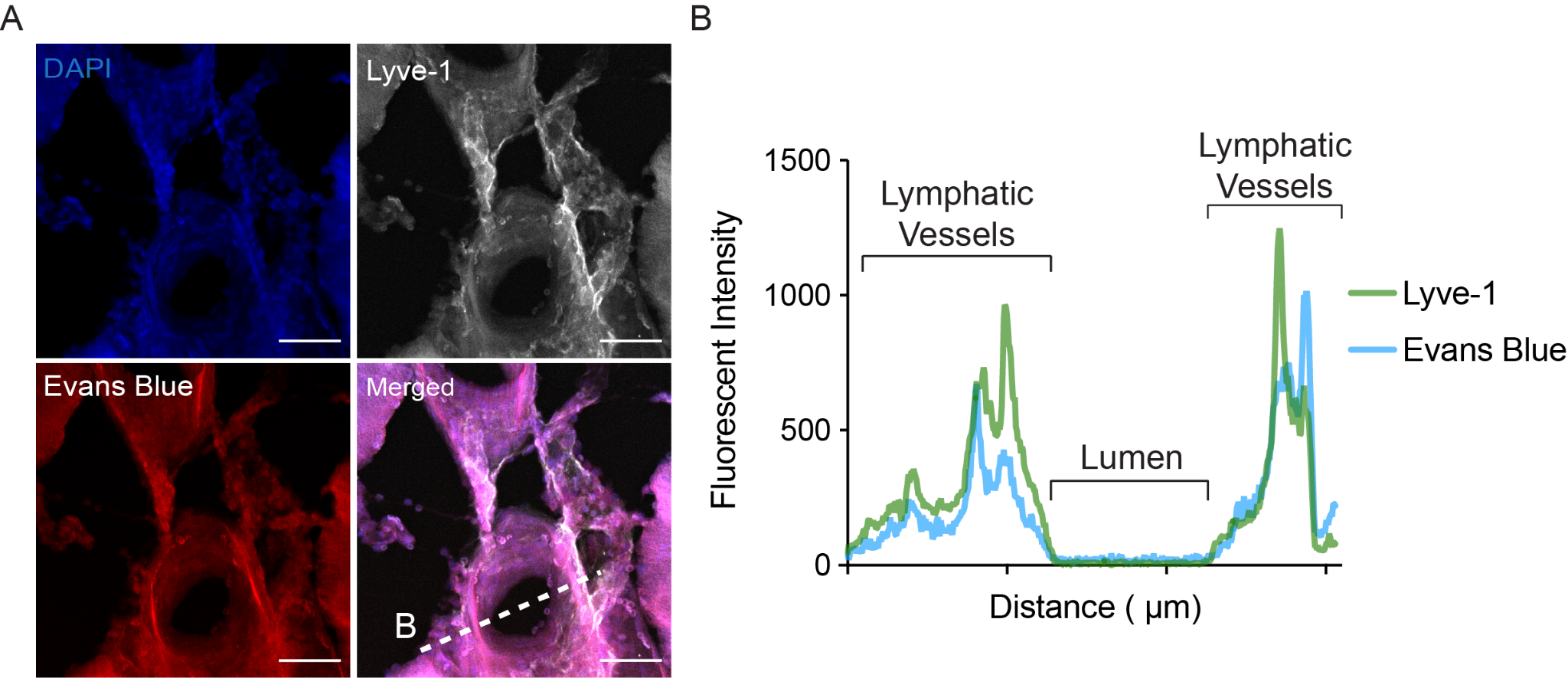

**Supplementary Figure 3. Lymphangiogenic vessels in the cribriform plate can drain the CSF after tMCAO.**

**(A):** After 7 days of tMCAO, 10% Evans Blue dye was injected into cisterna magna to confirm CSF drainage through lymphatics near the cribriform plate. Representative confocal images show overlap of Evans Blue dye and Lyve-1. Scale bars = 50 µm.

**(B):** The intensity profile plot shows co-localization of Evans Blue dye within Lyve-1^+^ lymphatic endothelial cells.

**Supplementary Figure 4. MAZ51 improves early post stroke motor outcomes**

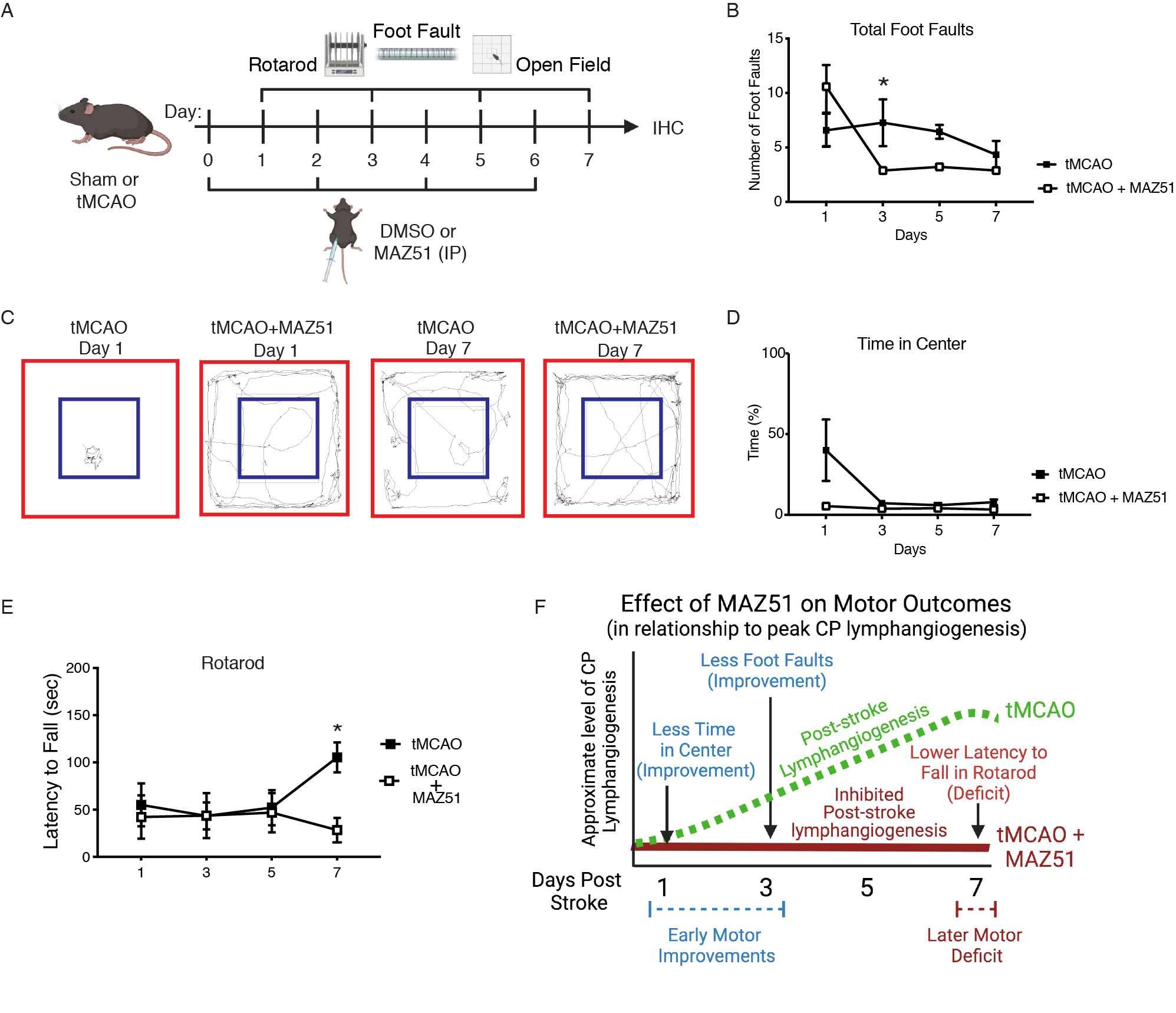

**Supplementary Figure 4. MAZ51 improves early post stroke motor outcomes**

**(A):** Ladder rung test was used to measure total number of foot faults to assess locomotor ability between control-treated tMCAO mice and MAZ51-treated tMCAO mice at day 1, 3, 5, and 7 (n=6 mice per group; mean ± SEM, *p < 0.05, two-way ANOVA)

**(B):** Percentage of time in center of the open-field box was measured between control-treated tMCAO mice and MAZ51-treated tMCAO mice at day 1, 3, 5, and 7 (n=6 mice per group; mean ± SEM, two-way ANOVA)

**(C):** Representative open-field track maps between tMCAO with control or MAZ51 at day 1 and 7 for comparison. Blue box indicates the center of the box and the red line indicates the peripheral boundary of the box. While control-treated tMCAO mice showed difficulties in movement, MAZ51-treated tMCAO mice showed improved locomotor activities at Day 1.

**(E):** Latency duration on rotarod was measured between control-treated tMCAO mice and MAZ51-treated tMCAO mice at day 1, 3, 5, and 7 (n=6 mice per group; mean ± SEM, *p < 0.05, two-way ANOVA)

**(F):** Timeline summary of motor function effects of MAZ51 treatment in relation to relative peak CP lymphangiogenesis (Day 7). Early improvements in motor recovery (Day 1-3), but late deficit in rotarod (Day 7) could be associated with sustained MAZ51 inhibition of lymphangiogenesis. Peak post-stroke lymphangiogenesis typically occurs at the CP on day 7 post stroke. Created using BioRender.com.

**Supplementary Figure 5. No changes to lymphatics after VEGF-C stimulation**

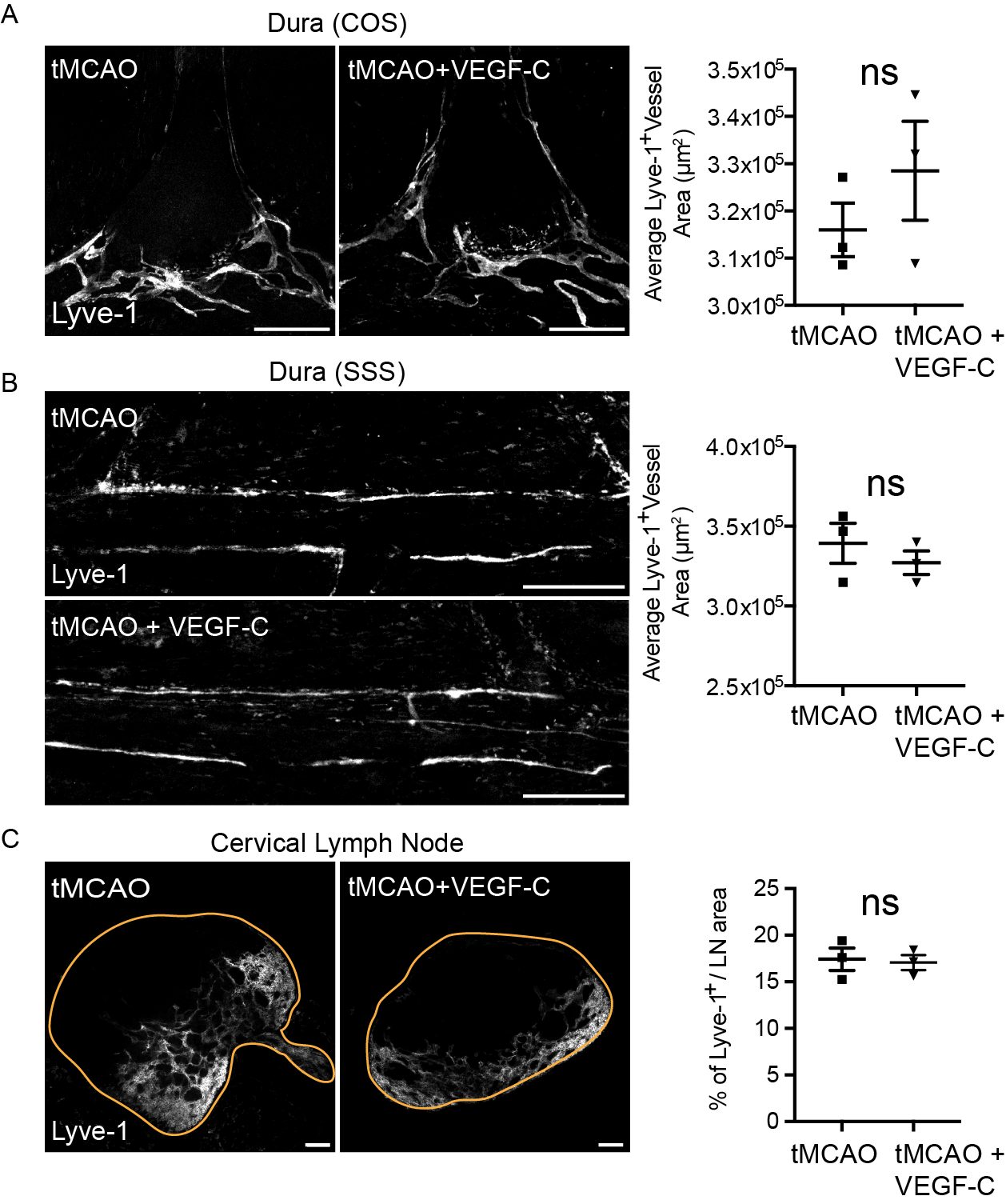

**Supplementary Figure 5. No changes to lymphatics after VEGF-C stimulation**

Coronal sections of COS of dural lymphatic vessels (**A**), SSS of dural lymphatic vessels (**B**) and sections of cervical lymph nodes (**C**) were stained with Lyve-1 fluorescent antibody after either control or VEGF-C156S treatment of tMCAO mice. Representative confocal images of lymphatic vessels near COS of dural lymphatics, cervical lymph nodes, and SSS of dural lymphatics. Scale bars = 500 µm for COS and SSS of dural lymphatic vessels, and 100 µm for cervical lymph nodes. Quantitation of each image (n=3 mice per group; mean ± SEM, unpaired Student’s t-test).
